# Design of a low-cost photomodulator for *in vivo* photoactivation of a mGluR5 inhibitor

**DOI:** 10.1101/2023.10.07.561310

**Authors:** Hans C. Ajieren, Andrew G. Fox, Ethan N. Biggs, Gabriel O. Albors, Amadeu Llebaria, Pedro P. Irazoqui

## Abstract

Severe side effects prevent the utilization of otherwise promising drugs in treatments. These side effects arise when drugs affect untargeted tissues due to poor target specificity. In photopharmacology, light controls the timing and the location of drug delivery, improving treatment specificity and pharmacokinetic control. Photopharmaceuticals have not seen widespread adoption in part because researchers do not always have access to reliable and reproducible light delivery devices at prices which fit within the larger research budget. In this work, we present a customizable photomodulator for use in both wearable and implantable devices. For experimental validation of the photomodulator, we photolyse JF-NP-26 in rats, producing raseglurant, a mGluR5 inhibitor shown to have antinociceptive effects in animal models. We show our photomodulator produces a significant reduction in pain response in the formalin model by photoreleasing raseglurant, indicating our photomodulator can successfully drive *in vivo* photopharmacology. We demonstrate modifications which enable the photomodulator to operate wirelessly. By documenting our photomodulator development, we hope to introduce researchers to a simple solution which significantly lowers the engineering barriers to photopharmacology research.

## Introduction

When administering or prescribing drugs, medical practitioners must balance treating a medical condition and maintaining normal function throughout the body. Many common drug delivery techniques result in systemic drug availability. Systemic drug availability can cause harmful side effects in untargeted tissues. Medical practitioners often limit the drug doses they prescribe to avoid these side effects. However, limiting drug doses to avoid side effects can limit the efficacy of drug treatments. Selective drug delivery techniques seek to overcome this limitation of systemic drug delivery.

Scientists utilize various techniques to deliver drugs more selectively: packaging drugs in hydrogels [1], repurposing extracellular vesicles [2], combining drugs with magnetic nanoparticles [3], [4], and modulating drug activity using light [5]–[12]. This last technique, photopharmacology, requires both light-responsive drugs and light-producing devices to control the light-responsive drugs. We use the term photomodulator to signify a device producing light to enable photopharmacology or other light-based biomedical interventions.

The growth of optogenetics as a research area has led to more equipment options for researchers employing light in biological experiments. Major vendors such as A-M Systems, Plexon, and Thorlabs have optogenetics kits in their catalogs, and more specialized companies like Neurolux and Teleopto offer their own optogenetics tools. Researchers have demonstrated innovative light systems for optogenetics [13]–[22]. These systems enable photopharmacology experiments, but they often include design choices which increase the per-unit cost of a system (like microfluidics for drug delivery) or prevent easy adaptation of the system to other applications. Commercial systems often prove quite expensive for exploratory research use, and the systems outlined in research papers often require specialized fabrication techniques like photolithography or micromachining. A low-cost, easily customizable photomodulator would facilitate photopharmacology research, especially as chemists develop new compounds to act on different anatomical targets.

With this motivation, we present our photomodulator design. To make our photomodulator easily reproducible, we designed the photomodulator circuitry using mass-produced integrated circuits (ICs) and light-emitting diodes (LEDs). We demonstrate the efficacy of this photomodulator by using it to photorelease raseglurant, a mGluR5 inhibitor shown to have an antinociceptive effect when released in the ventral posteromedial nucleus (VPM) of rodents subjected to the formalin pain model. We show how minor circuit modifications and inexpensively fabricated mechanical and optical components can adapt the photomodulator to different applications. We demonstrate how researchers can reconfigure the photomodulator to operate wirelessly and interface with a microcontroller for controls integration. The systems we develop demonstrate how researchers can build photomodulators at prices significantly below those quoted for commercial optogenetics systems.

## Results

### Photomodulator design and radiant power characterization

We began designing our photomodulator by exploring electronic methods for producing controllable amounts of light at a specified wavelength. We quickly selected light-emitting diodes (LEDs) as our light source. Many features make LEDs more suited for use in photomodulators than other light sources: low cost, small size, power efficiency, and availability in various wavelengths. Laser diodes share many of these features with LEDs, but LEDs tend to be less expensive than laser diodes.

Having chosen LEDs as our light sources, we needed to find a method for controlling the light produced by an LED. Selecting the right method for LED control comes down to understanding how an LED produces light. An LED produces light due to carrier recombination in its depletion region. This means an LED produces light in proportion to the current passing through the LED. The Shockley diode equation, included as Equation 1 in this paper, details the impact changes in voltage (*V*_*D*_) and temperature (*T*) have on the current (*I)* passing through a diode and explains why maintaining a constant current can produce more stable radiant power output from the LED than a constant voltage can.

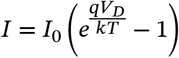

An LED driver is a specialized power source used to bias LEDs. Many LED drivers utilize current feedback to control the operating voltage. Our analysis of the Shockley diode equation explains the value of current feedback in LED control. IC manufacturers produce LED drivers at various levels of complexity. We selected the TPS61165 as the LED driver for our system to meet our goals of minimizing size and cost while maintaining control flexibility for future applications. The TPS61165, a six-pin IC, has a small footprint for an LED driver with current feedback and requires few external components, enabling minimization of the overall circuit footprint. We found LED drivers with greater efficiency (like the LT3791) and more control options (like the MAX1583), but they required more components, area, or power. We decided the TPS61165 met our size and functionality needs better than the other LED drivers we considered.

We found input voltage instability significantly impacted the operational consistency of the TPS61165. To address this, we added a voltage regulator to the photomodulator. We chose the ADP150, a four-pin voltage regulator IC. As with selecting our LED driver, we sought to minimize size and cost in selecting our voltage regulator. Employing a voltage regulator in our design simplified the later work of adapting the photomodulator to operate off different power sources. We added a Zener diode to the input of the voltage regulator to further protect the circuit from excessive input power.

In the tethered version of our photomodulator, we implemented a circuit for generating a pulse width modulation (PWM) signal with a hardware-controlled duty cycle. This PWM signal enabled us to modulate the LED driver output current from zero to its hardware-restricted maximum. For this circuit, we utilized an NE555, an inexpensive and well-documented timer IC. The NE555 enabled us to control the duty cycle of the PWM signal using a potentiometer.

Once we identified the ICs for our photomodulator circuit and selected our LEDs, we selected passive components based on guidelines in the IC datasheets and laid out a circuit board using circuit board design software. Figure 1 shows a schematic of the photomodulator circuit. We manufactured early versions of the photomodulator circuit board using a milling machine. Once we finalized the circuit board layout, we ordered PCBs through OSH Park. We manually populated the PCBs with the necessary circuit components.

**Figure 1:**
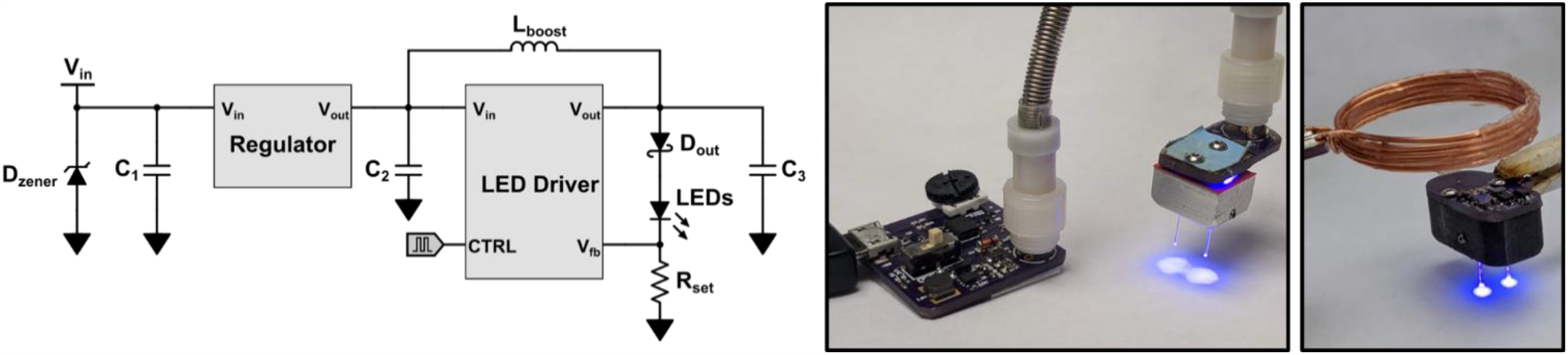
Photomodulator circuit diagram and photos. (Left) This simplified circuit schematic shows the essential photomodulator circuitry: a regulator, an LED driver, and supporting passives and diodes. A pulse-width modulation applied to the control (CTRL) pin of the LED driver modulates the photomodulator output current. (Center) A picture of the tethered version of the photomodulator. (Right) A picture of the wireless version of the photomodulator.

We began characterizing our photomodulator by assessing the linearity of radiant power control. We utilized two methods to control the radiant power output by the photomodulator: changing the set resistor and changing the PWM duty cycle. Figure 2 shows how changing these two controls impacted the radiant power output by the photomodulator. The output radiant power shows a strong linear relationship to both control variables (R^2^ = 0.999).

**Figure 2:**
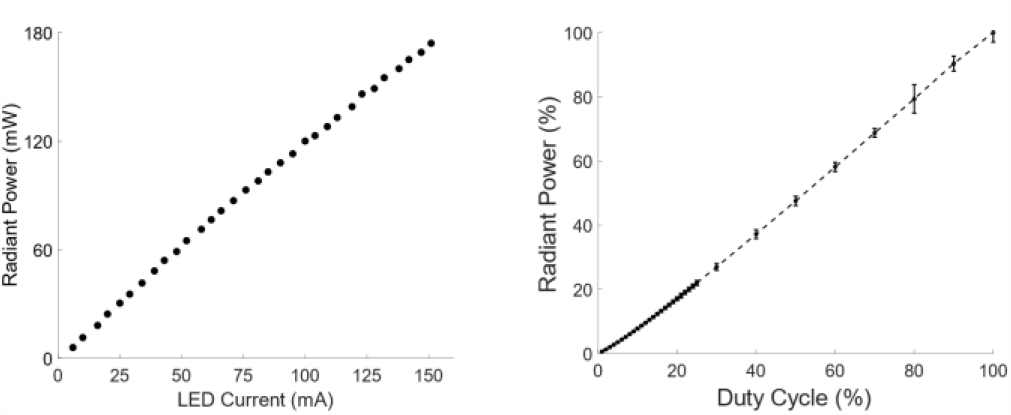
Photomodulator radiant power characterization. (Right) There is a strong linear relationship between photomodulator output current and LED radiant power (R^2^ = 0.99). (Left) When controlling the photomodulator output current by changing the duty cycle of the PWM control signal, the output radiant power responds linearly as well (R^2^ = 0.99).

### Behavioral photomodulation study

Having confirmed our ability to linearly control the radiant power output by the photomodulator, we moved to validating photomodulator functionality in an *in vivo* behavioral study. We modeled this study off the procedure Font and colleagues used to originally validate JF-NP-26 following their synthesis of the compound [12] and referenced Gong and colleagues for further detail [23].

Table 1 lists the behavioral experiment trial types. These trial types provide the statistical stratification needed to assess the necessity of both the drug and the light in photopharmacology. We performed 6 control trials, 4 drug-only trials, 5 light-only trials, and 4 treatment trials.

**Table 1:** Behavioral experimentation trial types. Each experimental trial type involved administering a different combination of intraperitoneal (IP) injection and photomodulation.

| Trial Type | IP Injection | Photomodulation<sup>[a]</sup> |
| --- | --- | --- |
| Control | Vehicle<sup>[b]</sup> | No |
| Drug-Only | JF-NP-26<sup>[c]</sup> | No |
| Light-Only | Vehicle | Yes |
| Treatment | JF-NP-26 | Yes |

[a] 5 minutes of 405 nm light illuminating the ventral posteromedial nuclei. [b] 10 mL/kg of 20% DMSO/20% Tween80/60% saline. [c] 10 mg/kg JF-NP-26 in 10 mL/kg of 20% DMSO/20% Tween80/60% saline.

Figure 3 shows the mean number of pain responses for the four trial types across six five-minute bins to visualize the temporal trends followed by each trial type. Figure 3 shows that control, drug-only, and light-only trials exhibited pain responses throughout the observation period while treatment trials only exhibited pain responses in the first ten minutes of the observation period. The treatment trials showed significantly fewer pain responses than the control, drug-only, and light-only trials in multiple bins.

**Figure 3:**
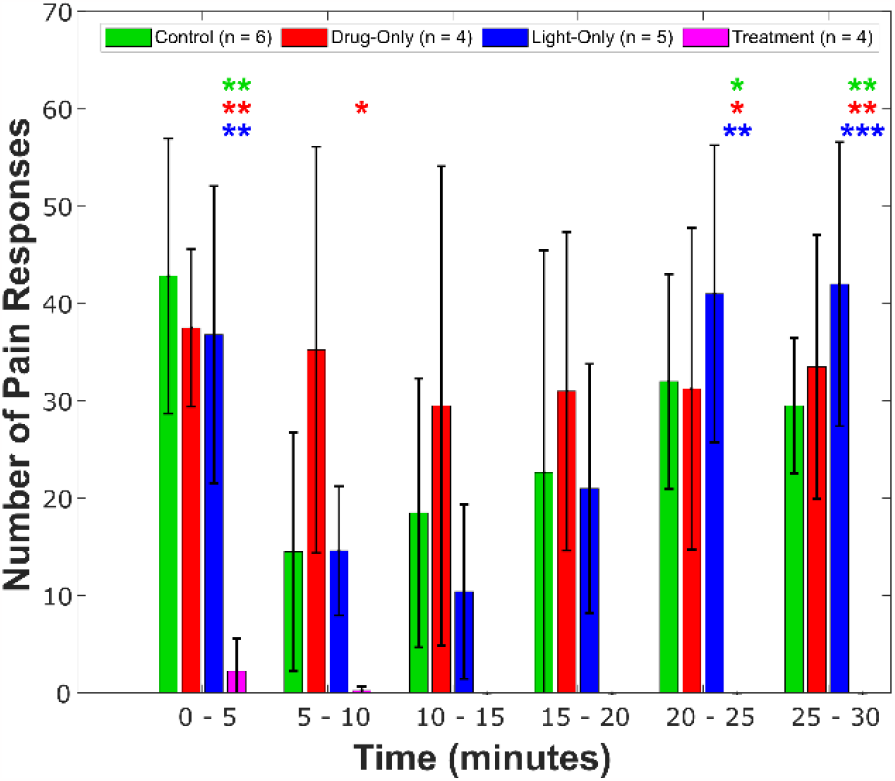
Time-binned pain responses. This histogram shows the mean number of pain responses recorded in five-minute bins for the different trial types. Error bars indicate standard deviation. While pain responses occur in every bin for control, drug-only, and light-only trials, no pain responses occur past the ten-minute mark in treatment trials. Asterisks indicate bins in which the mean pain response for the treatment group differs significantly from the mean pain response for another group. Asterisk color-coding matches bar color-coding. One, two, and three asterisks indicate p<0.05, p<0.01, and p<0.001, respectively. The mean pain responses of the other trail types were not statistically distinct from each other in any bin (p>0.05 for all other trial type combinations in all bins). For treatment trials, rats received JF-NP-26 and photomodulation. In drug-only trials, rats only received JF-NP-26, while in light-only trials, rats only received photomodulation. In control trials, rats did not receive JF-NP-26 nor photomodulation.

To assess the overall impact of treatment on pain response, we calculated the mean number of pain responses observed over the entire observation period for each trial type, as shown in Figure 4. One-way analysis of variance indicated that we had at least one statistically distinct trial group (p<0.001).

**Figure 4:**
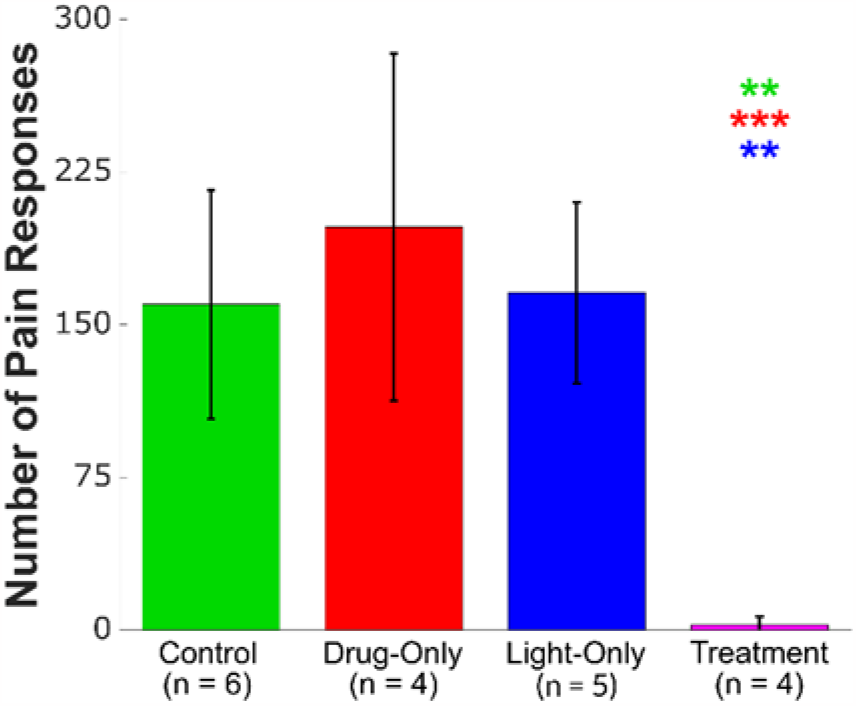
Mean pain responses. This graph shows the mean pain response of trials grouped by trial type with error bars indicating standard deviation. Asterisks indicate the significance with which the mean observed in the treatment trials differs from the means observed in the other groups. Asterisk color-coding matches bar color-coding. One, two, and three asterisks indicate p<0.05, p<0.01, and p<0.001, respectively. The mean pain responses of the other trail types were not statistically distinct from each other (p>0.05). For treatment trials, rats received JF-NP-26 and photomodulation. In drug-only trials, rats only received JF-NP-26, while in light-only trials, rats only received photomodulation. In control trials, rats did not receive JF-NP-26 nor photomodulation.

To find statistical differences between the trial groups, we conducted Tukey-Kramer multiple comparison tests. In control trials, we observed a mean pain response of 160.0±56.1 (the range indicates standard deviation). This mean pain response per trial was significantly affected by treatment (2.5±4.4, p=0.002) and not by drug-only (198.0±85.3, p=0.71) or light-only (165.8±44.5, p>0.99) intervention. This suggests the analgesic effect achieved in the treatment trials resulted from the photolysis of JF-NP-26 and not from the administration of JF-NP-26 alone nor from irradiation with 405 nm light alone. Figure 5 shows the chemical equation involved in this photolysis process.

**Figure 5:**
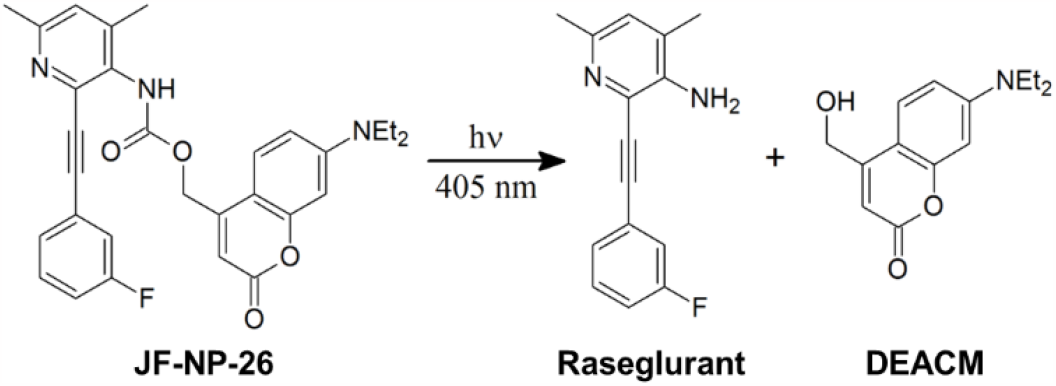
Photolysis of JF-NP-26. The photolysis of JF-NP-26 separates raseglurant (a mGluR5 inhibitor which acts as an antinociceptive drug) from DEACM (a photocaging side group).

### Wireless powering and system integration

Using a vector network analyzer, we measured the impedance of the receive coils fabricated on the wireless photomodulator PCBs across a frequency span of 500 kHz centered at 13.56 MHz, the center frequency of the ISM band we chose for wireless powering. We separated the impedance into its resistive (real) and reactive (imaginary) components and calculated the mean absolute percent error (MAPE) of these measurements using the following equation:

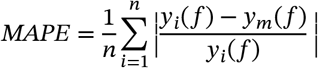

In this equation, *n* equals the total number of coils measured, *i* represents the index of the current coil, *y*_*i*_ equals the inductance or resistance of the current coil, and *y*_m_ equals the mean inductance or resistance. On average, the coils exhibited less error across the sampled frequencies for inductance (0.323%±0.010%) than for resistance (1.69%±0.43%). These average MAPEs fell below the typical 2% tolerance of the capacitors used in receive coil impedance matching. These MAPE calculations indicate the PCB manufacturing process yields highly consistent coils. Consistent coil impedance facilitates the reproduction of our wireless photomodulators; once you manage to impedance match the first receive coil with certain component values, you can use those values to effectively impedance match any coils with impedances close to the impedance of the first receive coil. Figure S1 shows a plot of the receive coil impedance measurements and calculated MAPEs against frequency.

Having measured the receive coil impedances, we assessed the performance of these coils in a wireless powering system. Figure 6 shows the uncoupled resonances of the coils and the power transfer efficiency between the coils at different distances. We calculated power transfer efficiency using the following equation (with results expressed as percentages):

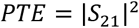

**Figure 6:**
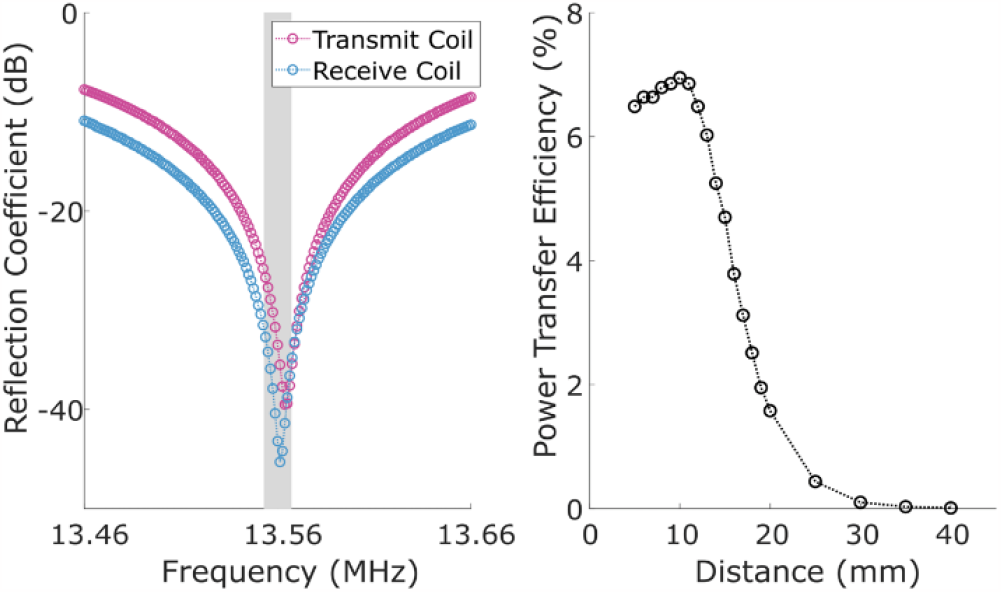
Resonance and power transfer efficiency of wireless powering coils. (Left) The unloaded resonances of the transmit coil and the receive coil fall within the 13.56 MHz ISM band (shaded region). (Right) The power transfer efficiency between the receive coil and the transmit coil initially increases slightly as coil separation increases to 10 mm and drops off steadily as separation increases past 10 mm.

Our results suggest we could further optimize the wireless powering system used with the wireless photomodulator. When designing the wireless photomodulator, we sought to maximize the receive coil inductance by maximizing the number of coil windings we fit into a given circuit board area. Increasing the number of receive coil windings increases the parasitic resistance in the receive coil, and this increase in resistance can reduce wireless power transfer efficiency. Improving the receive coil design would involve optimizing the number of coil windings and the width of the coil windings for a given coil area. Electromagnetics modeling software could help in optimizing coils for wireless powering.

When implemented in freely-moving animals, wireless photomodulators face variable coupling conditions due to the receive coil changing position. To address this, researchers implement antenna arrays and resonant cavities to create larger zones of adequate coupling. Researchers also implement dynamic impedance matching systems to actively improve coupling [24]. Transmit-side solutions face fewer constraints than receive-side solutions since the receiver must be attached to an animal.

With the photomodulator operating off wireless power, we moved to integrate the photomodulator into a larger wireless device. Figure S2 shows the photomodulator operating as a module attached to a Bionode, a neural recording and neuromodulation device developed and documented by our research group in previous work [25]. A PWM signal produced by the microcontroller of the Bionode controls the photomodulator, setting its output radiant power level.

Table 2 shows how the material costs of the photomodulators presented in this work compare to the costs of commercial systems. All prices listed in the table represent the price for a single wavelength bilateral photomodulator. These prices show how relatively inexpensive a custom photomodulator can be relative to catalogued commercial systems.

**Table 2:**
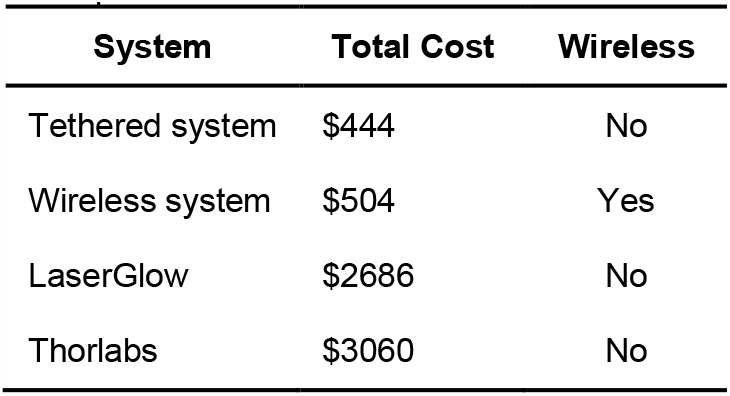
Photomodulator price comparison. The photomodulators developed in this work cost five times less than commercial systems capable of bilateral photomodulation.

| System | Total Cost | Wireless |
| --- | --- | --- |
| Tethered system | \$444 | No |
| Wireless system | \$504 | Yes |
| LaserGlow | \$2686 | No |
| Thorlabs | \$3060 | No |

## Conclusion

This study shows how commercially available circuit components and inexpensive fabrication techniques can produce a functional photomodulator. Access to an inexpensive photomodulator will benefit researchers synthesizing and validating new photopharmaceuticals. Lower photomodulator costs will enable greater budgeting efficiency in photopharmacology studies.

Recent electronics supply challenges emphasize the value of the generalized design of the photomodulator presented in this work. By using ICs with basic features throughout the photomodulator, we developed a photomodulator which facilitates the interchange of components with minimal circuit redesign. We leveraged this inter-changeability when faced with a backorder of the TPS61165 LED driver. We replaced the TPS61165 with another LED driver with an identical footprint and control scheme, allowing us to swap in this new LED driver without designing a new PCB.

The receive coil consistency achieved by designing the receive coil as part of the photomodulator PCB could greatly benefit researchers looking to use a wireless photomodulator without consistent access to a vector network analyzer. Impedance matching the first receive coil might take a long time, but subsequent impedance matching would only require repeating component values. Switching to a PCB coil design for the transmit coil would replicate this repeatable impedance matching on the transmit side of the system. Fabricating the coils for wireless powering on PCBs also saves researchers from having to learn time-consuming manual fabrication techniques.

This work showed a photomodulator working alongside a Bionode as an example of a system which could enable closed-loop photopharmacology. Pairing the photomodulator with a recording device enables photomodulation in response to heart rate, nerve activity, or some other biosignal. This would enable closed-loop photopharmacology experiments similar to those performed with electrical stimulation [26] or optogenetics [27].

Photopharmacology shows great promise for enabling novel methods to control drug activity. Researchers are particularly interested in photopharmaceuticals called photoswitches. A photoswitch switches between active and inactive isomeric forms with the application of light [28]. Multiple research teams have synthesized molecules that perform photoswitching [6], [10], [29]–[31]. Photoswitching compounds demonstrate photopharmacology’s greatest potential: precise control of drug activation and inactivation in both time and space. Minor adjustments to the photomodulator presented in this paper could produce a photomodulator capable of providing the different light wavelengths needed to fully control photoswitch activity. By facilitating the exploration of new photopharmaceuticals, our device should promote the growth of photopharmacology research and interventions.

## Acknowledgements

The authors thank Yvonne Chen, Brett Collar, and Trevor Meyer, for fabrication and procedural assistance and Ryan Budde, Brett Collar, and Dr. Jay Shah for providing manuscript feedback.

All applicable international, national, and/or institutional guidelines for the care and use of animals were followed. All procedures performed in studies involving animals were in accordance with the ethical standards of the institution or practice at which the studies were conducted (PACUC Protocol #1807001777).

## Author contributions

Conceptualization: HCA, AL, PPI

Data curation: HCA, AGF

Formal analysis: HCA

Funding acquisition: AL, PPI

Investigation: HCA, AGF, ENB

Methodology: HCA, ENB, AL, PPI

Project administration: GOA, PPI

Resources: GOA, PPI

Supervision: GOA

Validation: HCA, AGF

Visualization: HCA

Writing: HCA, AGF, ENB, GOA, AL, PPI

## Materials and Methods

### Circuit population

We manually populated the printed circuit boards (PCBs) used in this study. We used a soldering microscope with variable zoom to view the PCBs and a dental pick to distribute solder paste on pads before positioning components with tweezers. We used a reflow gun and a soldering iron to solder the components to the PCBs. This manual soldering technique allowed us to replace components and adjust component values as necessary without reflowing the entire PCB.

### Optode fabrication

We fabricated custom optodes for bilateral implantation in the ventral posteromedial nucleus (VPM). We obtained SFLC440 stainless steel ferrules and FP400URT optic fiber (optic fiber with a 400 μm core diameter and a numerical aperture of 0.50) from Thorlabs. We began optode fabrication by filing a notch near the flat end of a stainless steel ferrule. This notch would enable dental cement to anchor the optode during implantation. We used a Thorlabs S90R ruby fiber scribe and followed Thorlabs recommended procedure to obtain a 30 mm segment of uncoated optic fiber. We applied cyanoacrylate glue to the cladding at one end of the optic fiber before feeding this end into the flat end of the stainless steel ferrule. We fed the optic fiber through the ferrule until the end of the optic fiber slightly protruded from the tapered end of the ferrule. We then allowed the cyanoacrylate glue to cure, securing the fiber in the ferrule. With the fiber secured in the ferrule, we used the ruby fiber scribe to cut the fiber protruding from the flat end of the ferrule down to the appropriate length for implantation. Referring to the stereotaxic coordinates of the rat VPM documented by Paxinos and Watson [32]. With the optic fiber cut to the desired length, we polished the ends of the fiber following the polishing process recommended by Thorlabs. Figure S3 shows two optodes fabricated following this procedure. Custom fabrication allowed us to produce these optodes less expensively than if we bought optodes from a vendor like Thorlabs.

### Mechanical coupler design

With a photomodulator circuit constructed, we needed a tool to mechanically couple the photomodulator LEDs to fiber optic implants. Initially, we devised a bilateral coupler out of two Thorlabs ADAL3 quick-release interconnects. We saw an opportunity to improve the profile and mechanical integrity of our coupler by machining a custom coupler out of aluminum using a drill press. This machined aluminum coupler worked well for tethered experiments with the photomodulator but significantly reduced wireless powering efficiency when used with the wireless photomodulator. For this reason, we designed and 3D printed a polylactic acid (PLA) coupler for the wireless photomodulator. Figure S4 shows the different versions of the mechanical coupler used with our bilateral photomodulators.

### Radiant power measurement

We measured the radiant power produced by our bilateral photomodulator using the Thorlabs PM100D optical power meter. We performed the measurements in a room which allowed us to block out external light sources. We controlled the radiant power produced by the photomodulator both by changing the current setting resistor value and by delivering a PWM signal to the control pin of the LED driver.

### Coil Fabrication

Our wireless powering system involves two coils: a transmit coil and a receive coil. We fabricated a transmit coil by winding 18 AWG insulated magnet wire around a 32 mm diameter rod to create five congruent stacked windings. We used hot glue and cyanoacrylate glue to adhere the loops to each other and maintain the coil shape. We chose this size for our transmit coil because we wanted a mechanically robust coil which facilitated coil alignment for wireless powering. Smaller coils made with thinner magnet wire would deform more easily if dropped or mishandled, and larger coils would present more opportunities for misalignment during wireless powering.

We designed our receive coil as part of the circuit board for the wireless photomodulator to enable automated, reproducible receive coil fabrication. Manual coil fabrication would likely create a bottleneck when scaling photomodulator production for studies with a need for many photomodulators. We wanted a receive coil which matched the scale of the photomodulator circuit. This led us to set the outer diameter of the receive coil at 1 cm. With the manufacturing constraints of our PCB manufacturer, we managed to fit 12 concentric windings within thisdiameter. We took advantage of the double-sided fabrication of the PCB to add another set of 12 concentric windings to the coil, resulting in 24 total windings. All the receive coil windings curled in the same direction. This keeps current traveling in the same direction throughout the receive coil, leading to constructive interference and increased inductance. Figure S5 includes a picture of a transmit coil and a picture of a receive coil on a wireless photomodulator PCB.

### Impedance Matching

We used the same impedance matching strategy for transmit and receive coils. We utilized a vector network analyzer (VNA) to measure the impedance of a coil at 13.56 MHz. With this initial impedance measurement, we could determine the capacitor values needed to match the coil impedance to 50 Ω using the capacitive L network shown in Figure S5. We soldered capacitors in place to achieve the nominal values indicated by the equations and measured the resulting impedance of the impedance-matched coil. We added or subtracted capacitance as necessary to achieve an S11 magnitude less than or equal to -30 dB, a constraint which corresponds to an impedance mismatch of about 3% between the coil impedance seen through the matching network and the target impedance of 50 Ω.

### Surgical Procedure

A rat had ad libitum access to food and water until surgery. To begin the surgical procedure, we anesthetized the rat with 5% isoflurane in O2. A thermostatically controlled heating pad maintained rectal temperature at 37°C (507223F, Harvard Apparatus). We mounted the rat in a stereotaxic frame with the bregma-lambda axis horizontal and maintained anesthesia with 2-4% isoflurane in O2. Referring to the stereotaxic coordinates for the rat brain presented in the work of Paxinos and Watson [23], we established target coordinates for the VPMs (3.5 mm caudal, 3.0 mm lateral to bregma, 6.0 mm below the cortical surface) and bilaterally inserted the optic fiber implants with their tips at those coordinates. Figure S3 includes pictures of a rat with implanted optodes to demonstrate the location of the implants. We housed the rats individually with ad libitum access to food and water after surgery.

### Drug Preparation

We obtained the JF-NP-26 used for this study from Tocris Bioscience. To prepare the JF-NP-26 solution administered during experiments, we dissolved JF-NP-26 in dimethyl sulfoxide (DMSO) before adding saline (0.9% sodium chloride injection, USP) and Tween80 (procured from Sigma Aldrich) to the solution to achieve the desired vehicle formulation (20% DMSO, 20% Tween80, and 60% saline). We used a vortex mixer to mix the solution and an ultrasonicator to degas the solution. To prevent undesired JF-NP-26 photolysis, we shielded the centrifuge tubes containing JF-NP-26 solutions with aluminum foil. We stored unused JF-NP-26 at -80°C.

### Formalin Pain Model

Figure S6 shows the procedural timeline we followed for the formalin pain model. We anesthetized the rat with isoflurane and injected JF-NP-26 (10 mg/kg IP) dissolved in vehicle or an equivalent volume of vehicle. We then placed the rat in the observation chamber and attached the photomodulator to the rat’s ferrules, after which the rat recovered from anesthesia. Figure S7 includes a diagram of the tethered photomodulator setup and a picture of a rat undergoing photomodulation in the observation chamber. In trials involving photomodulation, 15 minutes after the IP injection, we turned on the photomodulator for 5 minutes. Following photomodulation, we disconnected the photomodulator from the implanted ferrules.

Following photomodulation, we injected 50 μL of 5% formalin into the hind paw. After administering the formalin injection, we returned the rat to the observation chamber and began thirty minutes of observation, recording the number of pain responses observed during this time. A rat had to either return his injected paw to the floor or take a visible pause from licking the injected paw for a repeated behavior to count as an additional pain response.

At the end of the observation period, we returned the rat to his individual cage and continued to observe the rat until the rat exhibited no pain responses for five minutes. Rats underwent a maximum of four trials of the experiment and recovered for at least 24 hours between each trial to allow for inflammation reduction in the injected paw and full metabolism of any drugs in their system. We alternated which hind paw received the formalin injection for subsequent trials. We conducted control, light-only, and treatment trials in a randomized order. The drug-only trials were not randomized with the other trials because we conducted these trials after review of trial data from the other groups to further validate the treatment outcome. In total, we used ten male Sprague Dawley rats (218-386 grams; 7-12 weeks; Envigo) to conduct these experiments.

As we performed the formalin pain model procedure, we had to evaluate the quality of the injections performed during the procedure. Sometimes rats experienced diarrhea following the IP injection, and we interpreted this as a sign of poor absorption of the injected solution. When administering the intraplantar formalin injection, we sometimes observed the formalin solution leaking from the injection site, resulting in a smaller noxious stimulus to the plantar tissue. If either of these undesired injection outcomes occurred, we terminated the trial at that point and did not collect trial data. However, we still counted the terminated trials against the maximum number of trials a rat could undergo.

**Figure S1:**
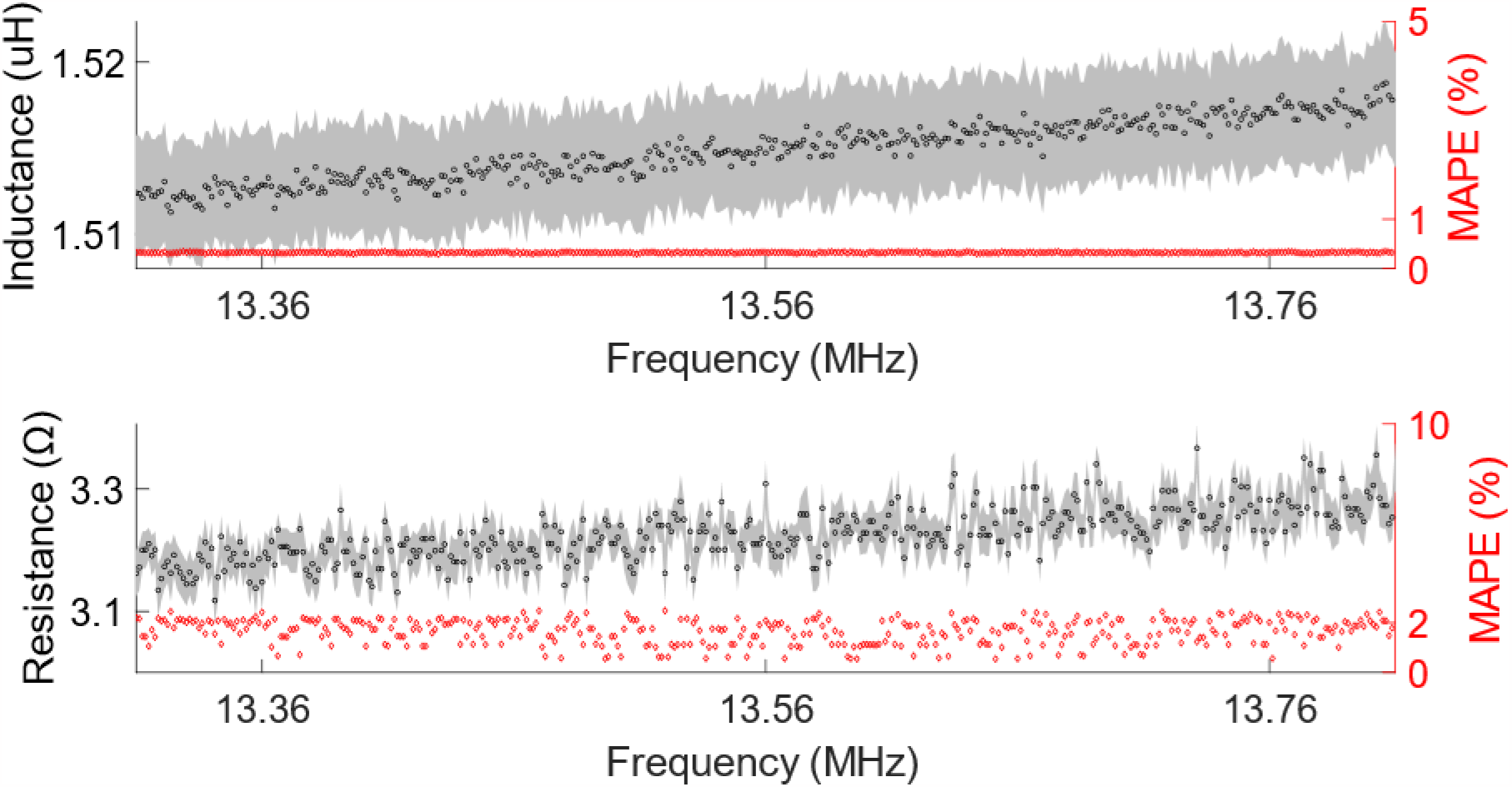
Wireless powering receive coil impedance measurements. (Top) Inductance and (bottom) resistance measurements of fifteen receive coils plotted against frequency. The shaded regions on the plots indicate the 90% confidence interval for the measurements. The datapoints in red correspond to the mean average percent error (MAPE) against frequency.

**Figure S2:**
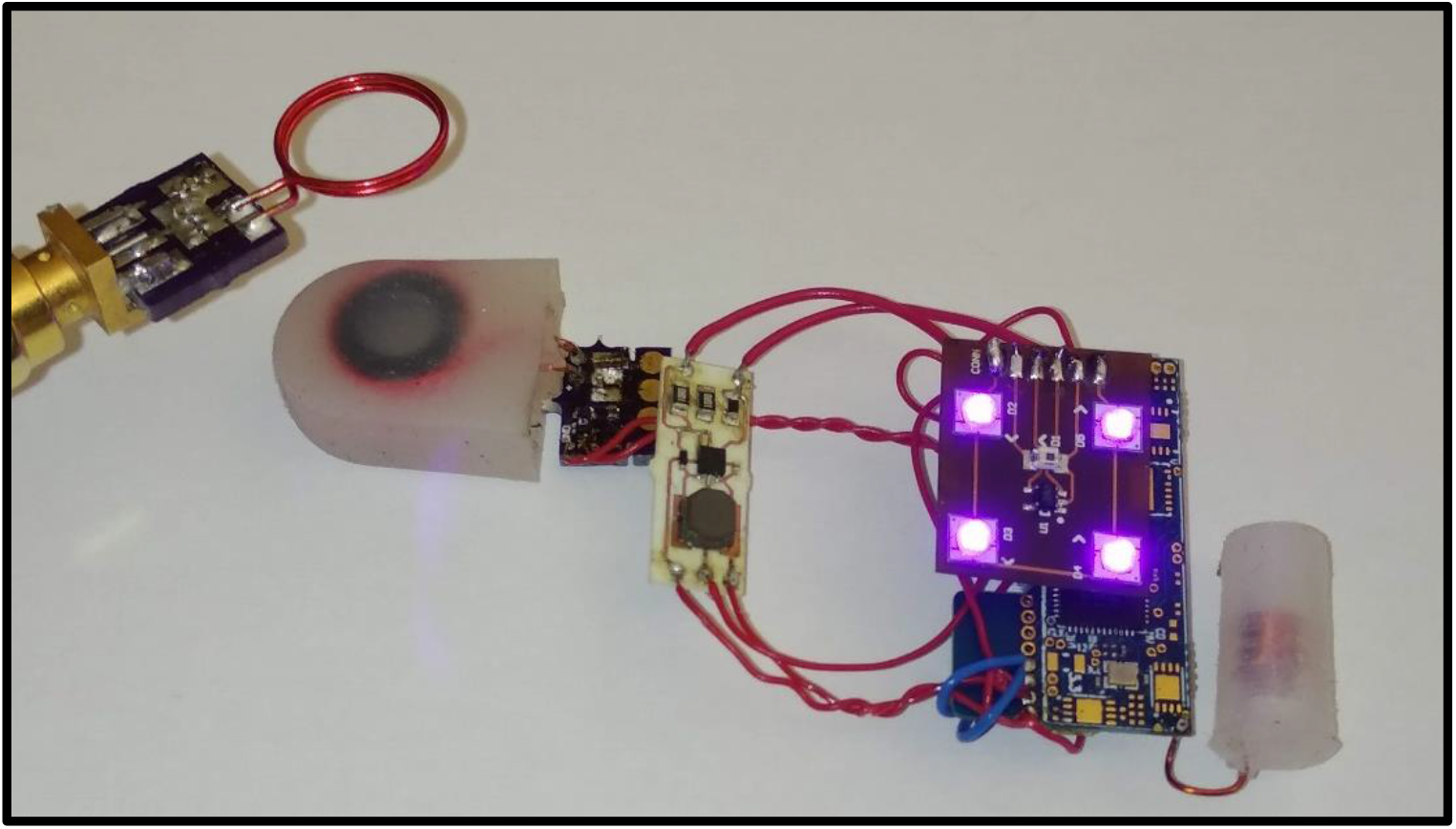
A photomodulator combined with a Bionode, a biosignal recording device. Connecting a photomodulator to a neuromodulation device such as a Bionode enables wireless control of the photomodulator in addition to wireless powering. Here, a Bionode (blue PCB) provides a PWM signal to the photomodulator (white PCB), controlling the radiant power output from the photomodulator LED array (brown PCB).

**Figure S3:**
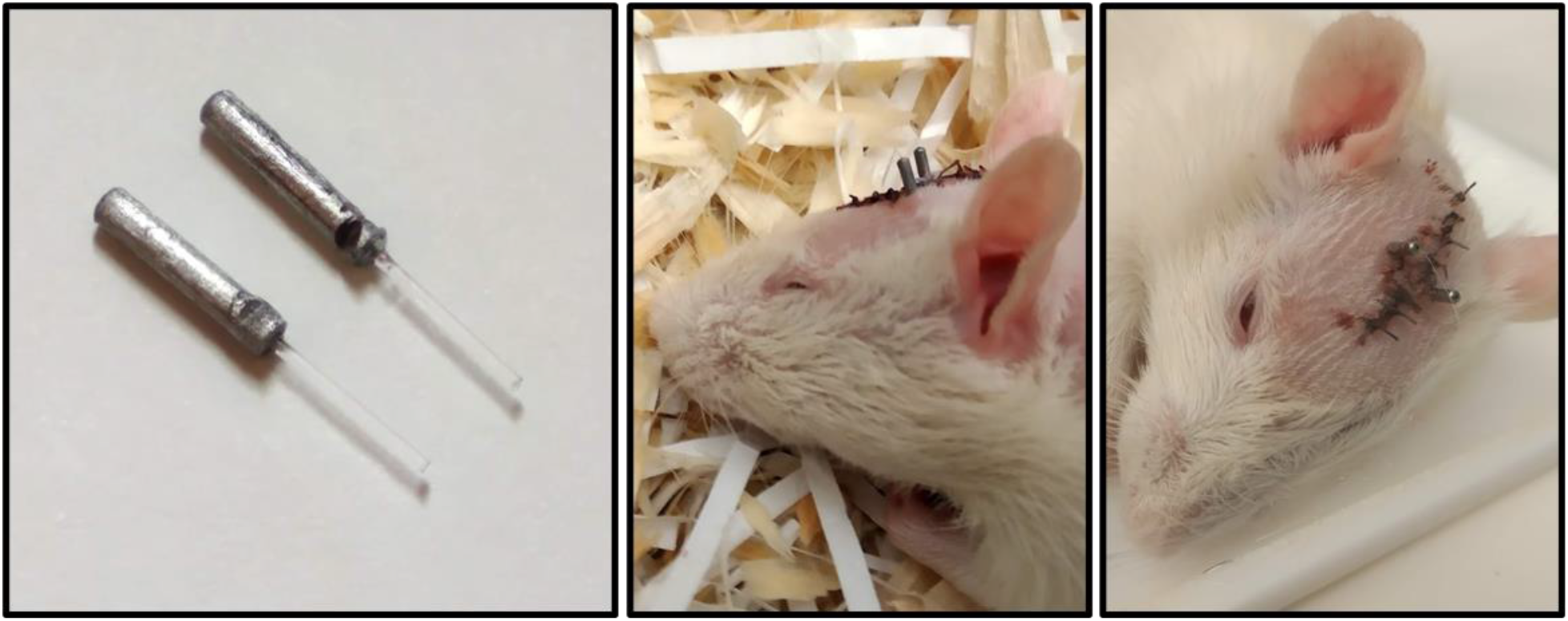
Fiber optic implants. (Left) Fiber optic implants. (Center and right) a rat after implantation of fiber optic implants, showing how ferrules remain exposed for interconnect attachment.

**Figure S4:**
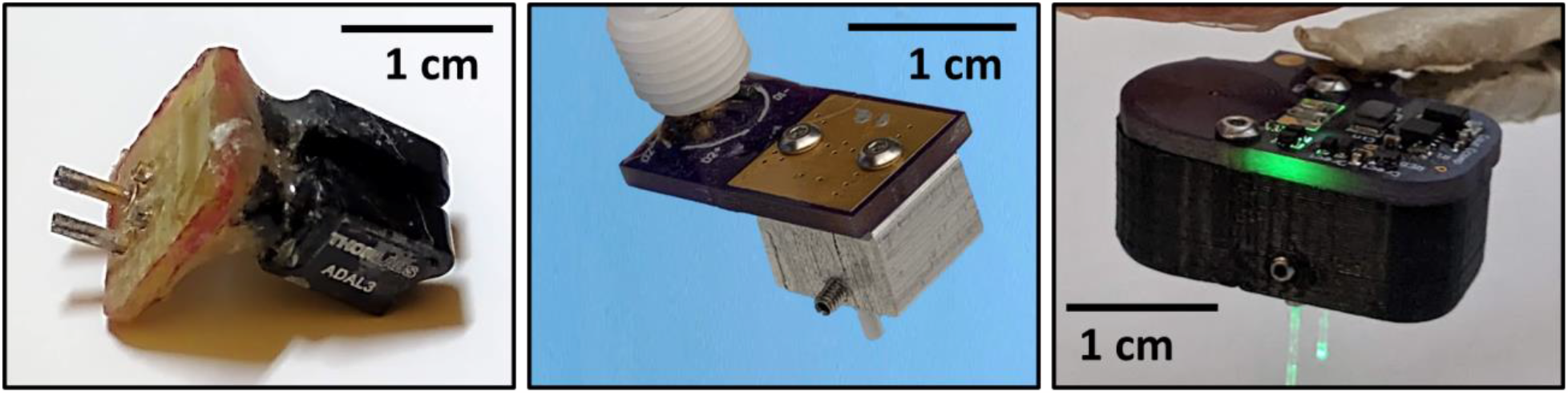
Mechanical coupler iterations. (Left) An LED module with a mechanical coupler formed using Thor Labs interconnects. (Center) An LED module with a mechanical coupler machined from aluminum. (Right) A wireless photomodulator with a 3D printed mechanical coupler.

**Figure S5:**
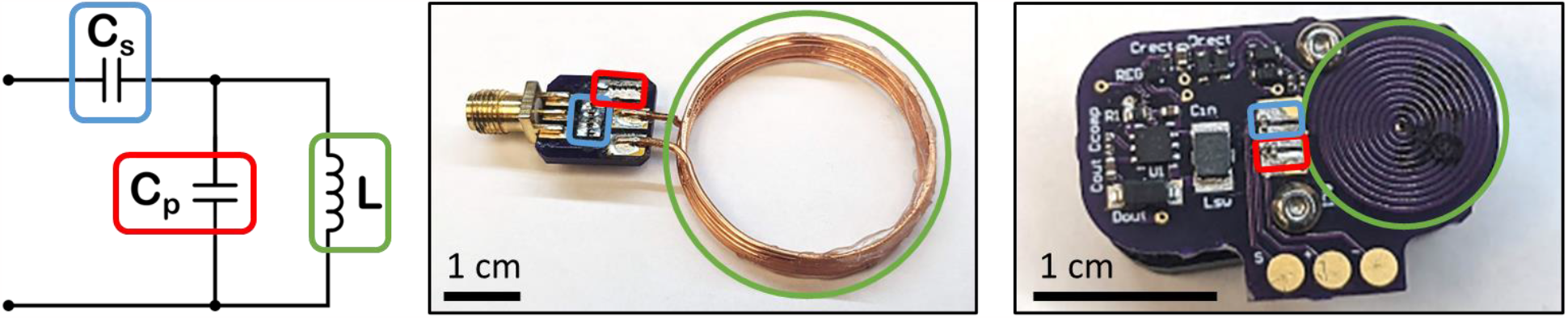
Wireless powering coils and matching networks. We used a two-capacitor L matching network (left) to match the impedances of the transmit coil (center) and the receive coil (right).

**Figure S6:**
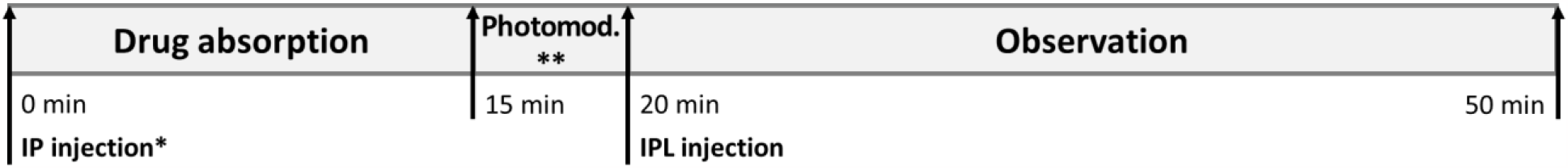
Procedural timeline for *in vivo* photomodulation experiments. A timeline for the behavioral experiments conducted to determine the efficacy of photomodulating JF-NP-26. Rats received different IP injections and photomodulation based on the trial type performed, as detailed in Table 1. All rats received the same intraplantar (IPL) injection of formalin to begin the observational period.

**Figure S7:**
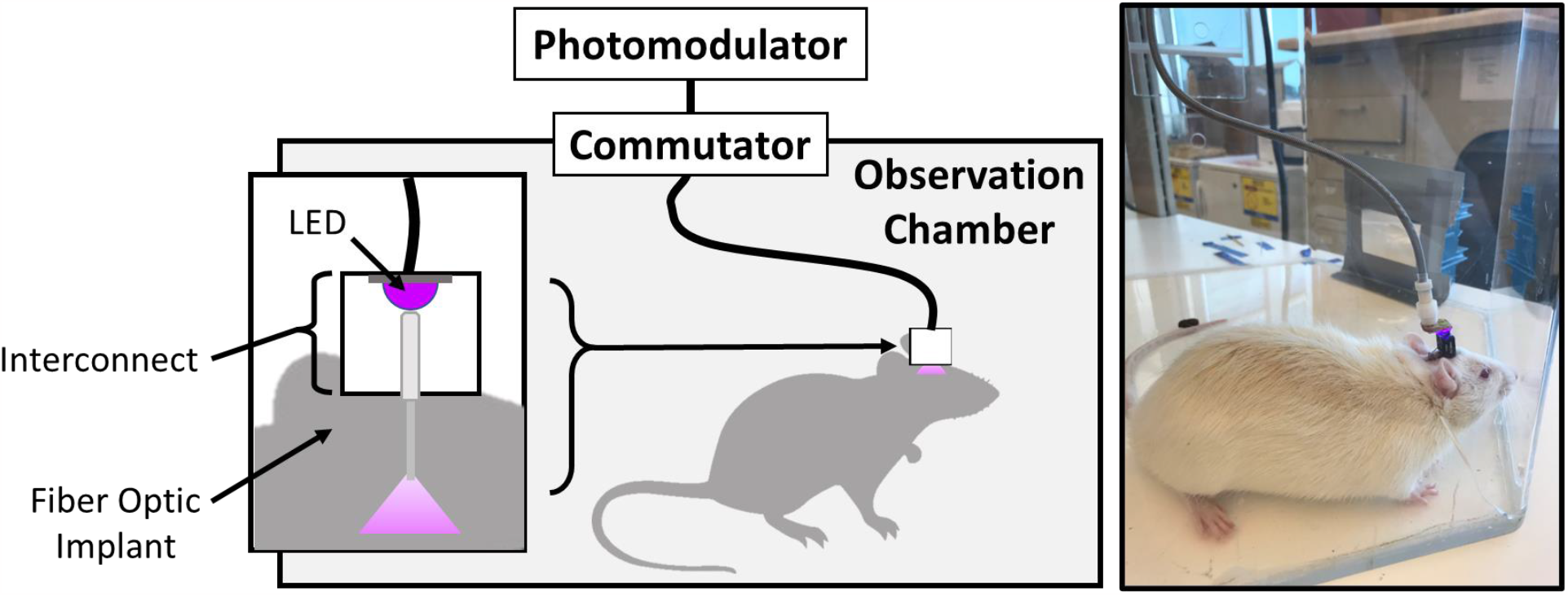
The behavioral observation chamber. This diagram and photograph show the chamber used for all behavioral experiments and the method used to mount LEDs to the implanted optic fibers for photomodulation. An interconnect couples the LEDs to the fiber optic implants, and a flexible cable connects the LEDs to a commutator, allowing the rat to move throughout the observation chamber. The commutator connects the LEDs to the photomodulator circuitry kept outside of the observation chamber.

